# A morphometric approach to the taxonomic dilemma of *Zonozoe drabowiensis* Barrande, 1872 and *Zonoscutum solum* Chlupáč, 1999 (Upper Ordovician, Czech Republic)

**DOI:** 10.1101/2025.09.11.675728

**Authors:** Lorenzo Lustri, Lukáš Laibl, Luis Collantes, Jana Bruthansová, Martina Nohejlová, Yu Liu, Stephen Pates

**Affiliations:** Yunnan Key Laboratory for Palaeobiology, Institute of Palaeontology, Yunnan University, 650091, Kunming, China; MEC International Joint Laboratory for Palaeobiology and Palaeoenvironment, Yunnan University, 650500 Kunming, China; Czech Academy of Sciences, Institute of Geology, Rozvojová 269, 165 00, Prague 6, Czech Republic; Department of Palaeontology, National Museum, Cirkusová 1740, 193 00, Prague 9, Czech Republic; Czech Geological Survey, Klárov 3, 11800, Prague, Czech Republic; School of Geography, Geology and the Environment, University of Leicester, Leicester, UK; Department of Earth Sciences, University College London, Gower Street, London, WC1E 6BT, United Kingdom; Department of Genetics, Evolution and Environment, University College London, Gower Street, London, WC1E 6BT, United Kingdom

**Keywords:** *Zonozoe drabowiensis*, *Zonoscutum solum*, elliptical Fourier analysis, Vicissicaudata, Artiopoda

## Abstract

*Zonozoe drabowiensis* Barrande, 1872 and *Zonoscutum solum* Chlupáč, 1999 are rare and incompletely preserved arthropods from the Upper Ordovician of the Czech Republic. Their classification has been a subject of debate for over a century due to the limited number of specimens, lack of knowledge related to post-cephalic morphology and the absence of clear diagnostic features. They were previously considered as members of Aglaspidida, an extinct group of arthropods from the Cambrian and Ordovician, within Vicissicaudata, a branch of the larger arthropod clade Artiopoda. Herein, we analysed the cephalic outlines of *Zonozoe* and *Zonoscutum* to determine whether their shapes align more closely with vissicaudatans than with other early Palaeozoic arthropods, offering a new morphological perspective on their systematics. We assembled a data set of cephalic outlines each representing one of thirty-three early Palaeozoic species, including *Zonozoe*, *Zonoscutum*, nine euchelicerates, six aglaspidids, three cheloniellids, and a selection of other artiopodans. We quantified their shape using elliptical Fourier analysis. Principal component analysis (PCA) and linear discriminant analysis (LDA) place *Zonozoe* within the vicissicaudatan morphospace, and *Zonoscutum* in their proximity, clearly distinguishing them from euchelicerates. These data add support to the most conservative classification of *Zonozoe* and *Zonoscutum* within Artiopoda, while strengthening the case for a more specific affinity with Vicissicaudata, helping to resolve a 150-year-old taxonomic uncertainty. More broadly, this study demonstrates the value of outline-based morphometrics in aiding systematic hypotheses when discrete characters are unavailable or scarce, offering a reproducible tool for re-evaluating other problematic fossils.

## INTRODUCTION

*Zonozoe drabowiensis* Barrande, 1872 and *Zonoscutum solum* are enigmatic arthropods known from the Letná and the Libeň Formations of the upper Ordovician of the Czech Republic (Chlupáč, 1965, 1999b; Rak et al., 2009; Vokáč & Grigar, 2024). Barrande (1872) originally described *Zonozoe* as an arthropod, specifically as a large carapace of an ostracod. Later on, Novák (1887) in his unpublished manuscript considered *Zonozoe* to be a merostomate arthropod close to Synziphosurina (Chelicerata) (Chlupáč, 1963, 1965, 1999b; Rak, 2009). Since then, these fossils have not received much attention until the second half of the twentieth century. Chlupáč (1963) assigned *Zonozoe* to the group Aglaspidida (Walcott, 1911) – a group of extinct marine arthropods, known from the Cambrian and Ordovician strata. However, this assignment remains uncertain (Chlupáč, 1963). It should also be noted that at that time, Aglaspidida was considered as one of the basal-most groups within Xiphosura (Chelicerata). Subsequent studies have revisited the taxon, reiterating its general association with Aglaspidida and/or the Merostomata, but always with a low degree of confidence (Chlupáč, 1965; Hou, 1997; Chlupáč, 1999a; Rak et al., 2009). As for *Zonoscutum*, the only fossil ascribed to this species was described in Chlupáč (1999a), who tentatively assigned it to Aglaspidida.

During the one hundred and fifty years since the discovery of *Zonozoe*, and twenty-five years for *Zonoscutum,* their classification has been complicated by two main factors. Firstly, the material ascribed to both *Zonozoe* and *Zonoscutum* is scarce, incomplete and not well preserved (Ortega□ernández et al., 2013; Van Roy et al., 2022). There are currently only eight known specimens of *Zonozoe* and only one specimen of *Zonoscutum,* all of which are isolated moulds of cephala (Chlupáč, 1999b; Rak et al., 2009; Vokáč & Grigar, 2024), preserved in quartzose sandstones of the Libeň and Letná Formations. In most cases, the specimens preserve only the general vaulted cephalon outline with a raised longitudinal region terminated with a ridge bearing what is a vague indication of the eyes (Rak et al., 2009; Van Roy et al., 2022). The only exception is a single specimen of *Zonozoe* preserving what has been referred to as glabellar furrows (Rak et al., 2009) and a specimen of *Zonoscutum* preserving similar structures but in a different position (Chlupáč, 1999a). Secondly, the classification of *Zonozoe* and *Zonoscutum* has been further complicated by the complex history of the classification of aglaspidid arthropods themselves, especially during the second half of the twentieth century. Their overall dorsal morphology led several authors to consider the aglaspidids as an order of the subclass Xiphosura (Størmer, 1955), grouping Aglaspidida with Xiphosura and Eurypterida, as part of the Merostomata (Raasch, 1939; Størmer, 1944; Eldredge & Smith, 1974) or to consider Aglaspidida as the sister group to chelicerates (Weygoldt and Paulus, 1979). This is in contrast to more recent aglaspidid phylogenies, which emphasize ventral anatomical features as key to understanding their true affinities (Briggs et al., 1979; Hesselbo, 1992; Ortega□Hernández et al., 2013; Lerosey-Aubril, Paterson, et al., 2017; Lerosey-Aubril, Zhu, et al., 2017). Currently, they have been placed within the broader clade Artiopoda, as part of the clade Vicissicaudata (Ortega□Hernández et al., 2013; Lerosey-Aubril, Paterson, et al., 2017; Lerosey-Aubril, Zhu, et al., 2017). Until very recently, the absence of strong defining characteristics for the clade has made Aglaspidida prone to being used as a wastebasket taxon (Plotnick & Wagner, 2006). This could be the case for the poorly preserved *Zonozoe* and *Zonoscutum*, which lack strong anatomical evidence supporting placement within aglaspidids. Other factors that led previous workers to assign *Zonozoe* and *Zonoscutum* to Aglaspidida are likely not only the general similarity between their cephala, but also the known (or suspected) presence of aglaspidids in the same paleogeographic region (Chlupáč, 1963, 1965; Van Roy, 2005; Ortega-Hernandez et al., 2010) and the weak resemblance with other arthropod groups (e.g. position of the eyes and glabella if compared with trilobites).

Later authors commenting on *Zonozoe* and *Zonoscutum* decided to have a more parsimonious approach. Ortega□Hernández et al. (2013) opted for the exclusion of those taxa from an extensive phylogenetic analysis of Vicissicaudata due to the lack of compelling taxonomically relevant features. Van Roy et al. (2022), instead, revised *Zonozoe* and *Zonoscutum* to critically assess their possible affinity with *Triopus draboviensis* Barrande, 1872. In their re-examination of the material, the authors did more than simply dismiss the affinity with *Triopus*; they advanced a conservative and more parsimonious taxonomic classification for both *Zonozoe* and *Zonoscutum*. They suggested that, at most, the fossils could be broadly classified as Artiopoda, and argued that any attempt to refine the classification beyond that level would be unwarranted (Van Roy et al., 2022). Known only from cephalic moulds, both *Zonozoe* and *Zonoscotum* lack the diagnostic features of Vicissicaudata, all of which are post-cephalic (Lerosey-Aubril, Zhu, et al., 2017; Briggs et al., 2023). They also lack several defining traits of Aglaspidida, including all post-cephalic diagnostic characters, most notably the modified posterior appendages, as well as evidence of a biomineralized cuticle and dorsal ecdysial sutures (Ortega□Hernández et al., 2013; Lerosey-Aubril, Zhu, et al., 2017). As a result, they cannot be confidently classified through standard phylogenetic or comparative analysis (Ortega□Hernández et al., 2013; Van Roy et al., 2022), a conclusion fully supported by this study. While the enigmatic *Zonozoe* and *Zonoscutum* are unlikely to be key taxa in contributing to resolving the phylogeny of Aglaspidida, a more accurate assignment of this species to this group may yield valuable palaeoecological and paleobiogeographical insights.

In order to contribute to resolving the taxonomic dilemma surrounding *Zonozoe* and *Zonoscutum* more effectively, we adopted an alternative approach to phylogenetic analysis and classical comparative anatomy, focusing instead on an overlooked well preserved feature in these fossils: the overall cephalic outline. We compared the outline of the cephalon of *Zonozoe* and *Zonoscutum* with thirty-three other arthropods. We implemented a principal component analysis (PCA) and linear discriminant analysis (LDA) on the study of the cephalic outline to extract the maximum morphological information from the limited material of *Zonozoe* and *Zonoscutum*. This methodology has been used in paleobiology to compare the outline of different fossils and can provide information on their taxonomy, ecology, and development (Gevirtz, 1976; Healy-Williams & Williams, 1981; Burke et al., 1987; Foote, 1989; Pates et al., 2021; Ma et al., 2023; Braig et al., 2024; Drage & Pates, 2024; Collantes & Pates, in press). A significant advantage of this methodology over landmark-based approaches is that it does not rely on discrete morphological characters, facilitating quantification of shape variation when such discrete characters are absent. Combined with the classical taxonomic work carried out by various authors over the years (Chlupáč, 1963, 1965, 1999b; Rak, 2009), our approach offers a new line of evidence, contributing to either confirming or rejecting previous hypotheses about the taxonomic affinities of these fossils. Both *Zonozoe* and *Zonoscutum* plotted within the Vicissicaudata area of morphospace, and thus the shape of their cephala concurs with a vicissicaudate affinity.

## MATERIALS AND METHODS

### Species involved in the study and the criteria of selection

In addition to one *Zonozoe* and one *Zonoscutum* specimen (Figure 1), nine euchelicerates, two nektaspids, six non-vicissicaudatans artiopodans and fourteen vicissicaudatans arthropods (including six aglaspidids and three cheloinellids), representing thirty-three species, were included in our study. Trilobites were excluded for several reasons: 1) While Cambrian and/or Ordovician nektaspids, euchelicerates, and vicissicaudatans are morphologically rather conservative—at least in the general shape of their cephala/prosomas (cf. Fig. 1; Lerosey-Aubril et al., 2017; Pérez-Peris et al., 2021)—this is not the case for trilobites. The cephalic morphology of trilobites is extremely diverse, particularly during the Ordovician period (e.g., Drage & Pates, 2024; Balseiro et al., 2025), and such diversity would necessarily introduce substantial noise into our dataset. This study aims to build on previous taxonomic work and, given the limitations of our methodology, to avoid introducing additional variability that would likely reduce the predictability of our analyses; (2) *Zonozoe* and *Zonoscutum* have never been considered closely related to Trilobita; and (3) the position of the eyes and glabella in *Zonozoe* and *Zonoscutum* differs markedly from that of any contemporaneous trilobite.

**Figure 1.**
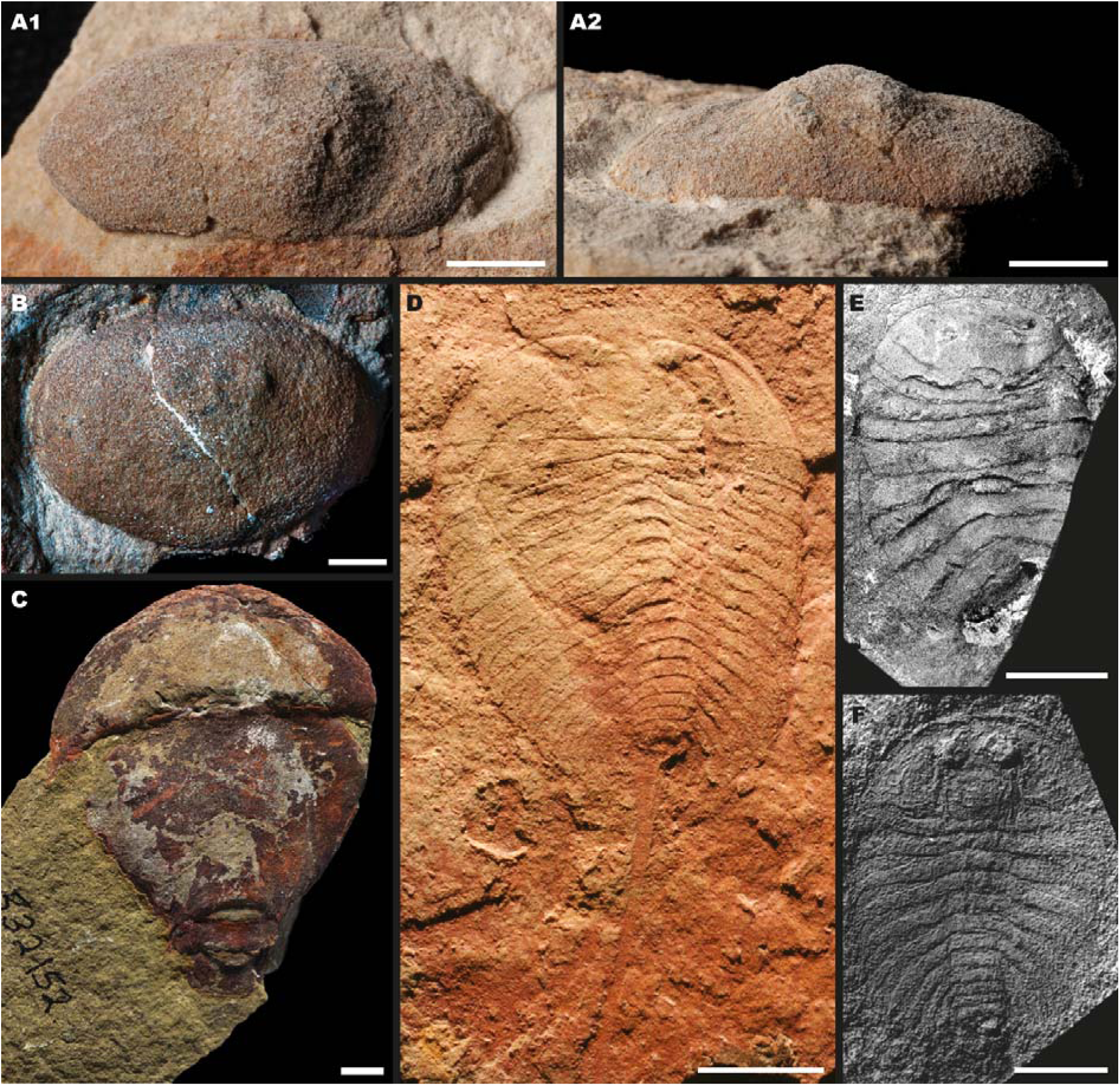
Representative cephala of *Zonozoe drabowiensis*, *Zonoscutum solum*, euchelicerate arthropods, vicissicaudatan cheloniellids, vicissicaudatan aglaspidids, and non-vicissicaudatan artiopodans featured in this study. A1, *Zonozoe drabowiensis*, dorsal view; A2, *Zonozoe drabowiensis*, frontal view; B, *Zonoscutum solum*, dorsal view, modified from Van Roy et al. (2022); C, Unnamed Xiphosura from the Fezouata biota, modified from (Lamsdell et al., 2013). D, *Eozetetes gemmelli*, modified from (Edgecombe et al., 2017); E, *Neostrabops martini*, modified from (Caster & Macke, 1952); F, *Aglaspis barrandei*, modified from (Hesselbo, 1992). Scale bar equals 5 mm.

**Figure 2.**
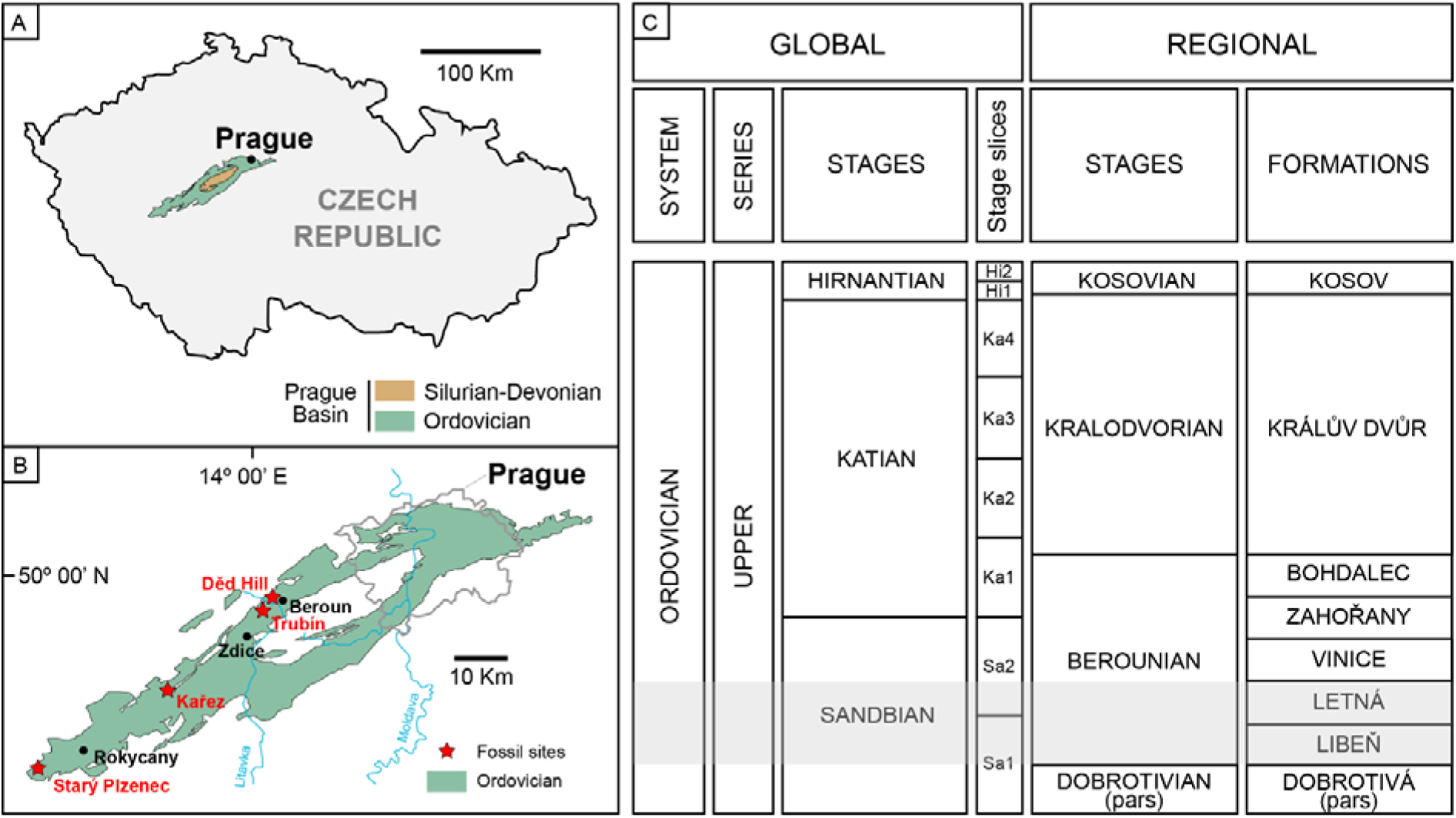
Geographical and stratigraphical position of the localities where *Zonozoe drabowiensis* and *Zonoscutum solum* were collected. A, Map of the Czech republic; B, Stratigraphical column of the upper Ordovician (modify after (Van Roy et al., 2022)).

The most complete and horizontally preserved specimen of *Zonozoe* (Lectotype NM L 23586) was chosen, while *Zonoscutum* is known from only one specimen (Holotype NM L 33021). The selection from the literature of other taxa, was made following four main criteria: 1) to include Early Palaeozoic diversity from various geographic regions, but with particular emphasis on species coeval and sympatric to *Zonozoe* and *Zonoscutum* (e.g. *Paraeurypterus anatoliensis* Lamsdell et al. 2013); 2) to exclude taxa with aberrant cephalic outlines; 3) to favour those potentially related to *Zonozoe* and *Zonoscutum* (e.g. *Eozetetes gemmelli* Edgecombe et al. 2017 and *Chlupacaris dubia* Van Roy, 2005); and 4) to maximize higher-level taxonomic diversity (e.g. *Carimersa neptuni* Briggs et al., 2023 and *Lunataspis borealis* Lamsdell et al., 2022). A summary of the species and specimens included in the analysis, along with inventory numbers, geological age, geographic provenance, and references to the original sources from which raw images were obtained, is provided in Table 1, while the specimen sample, one for each high taxonomic group, is shown in Figure 1.

**Table 1.**
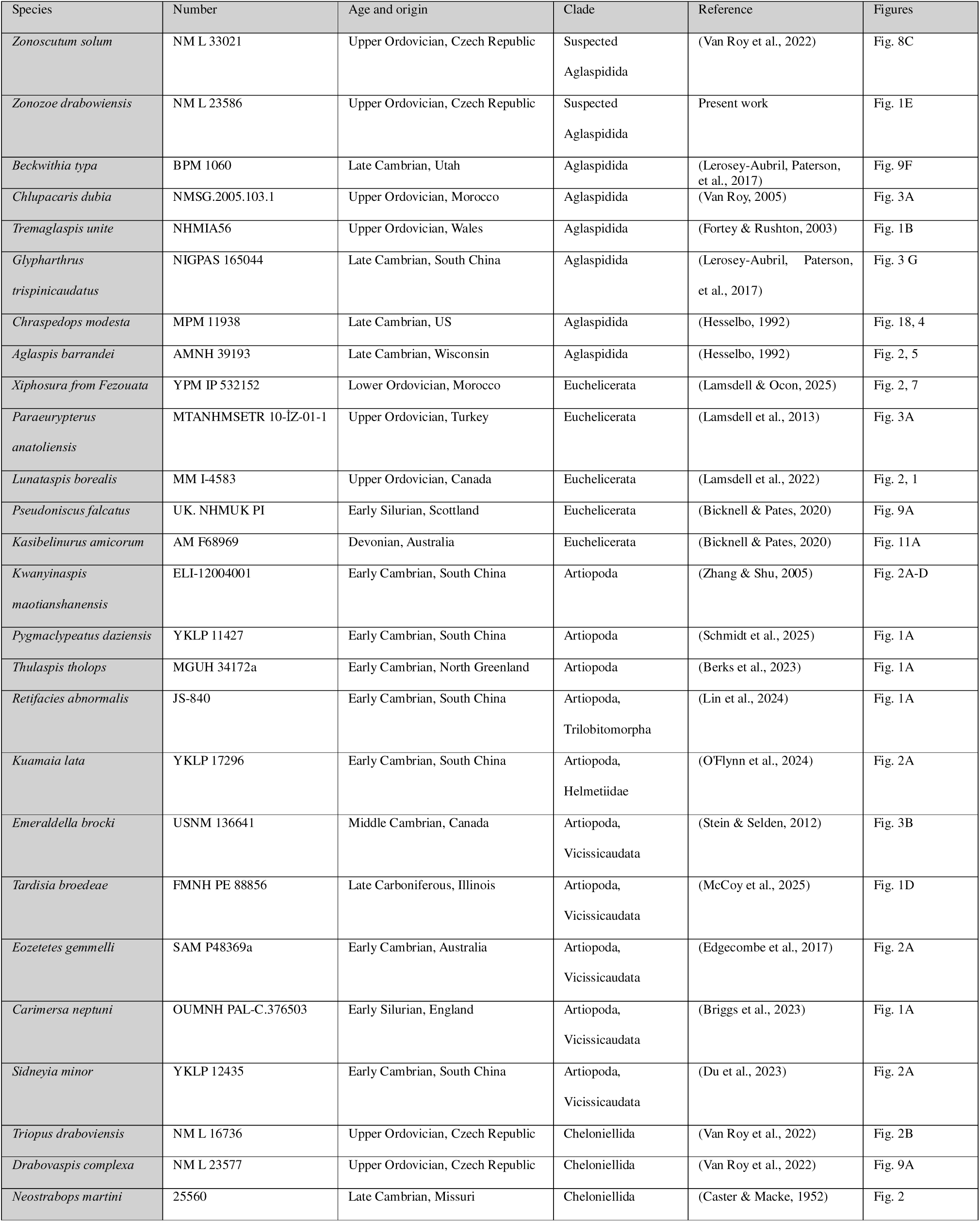
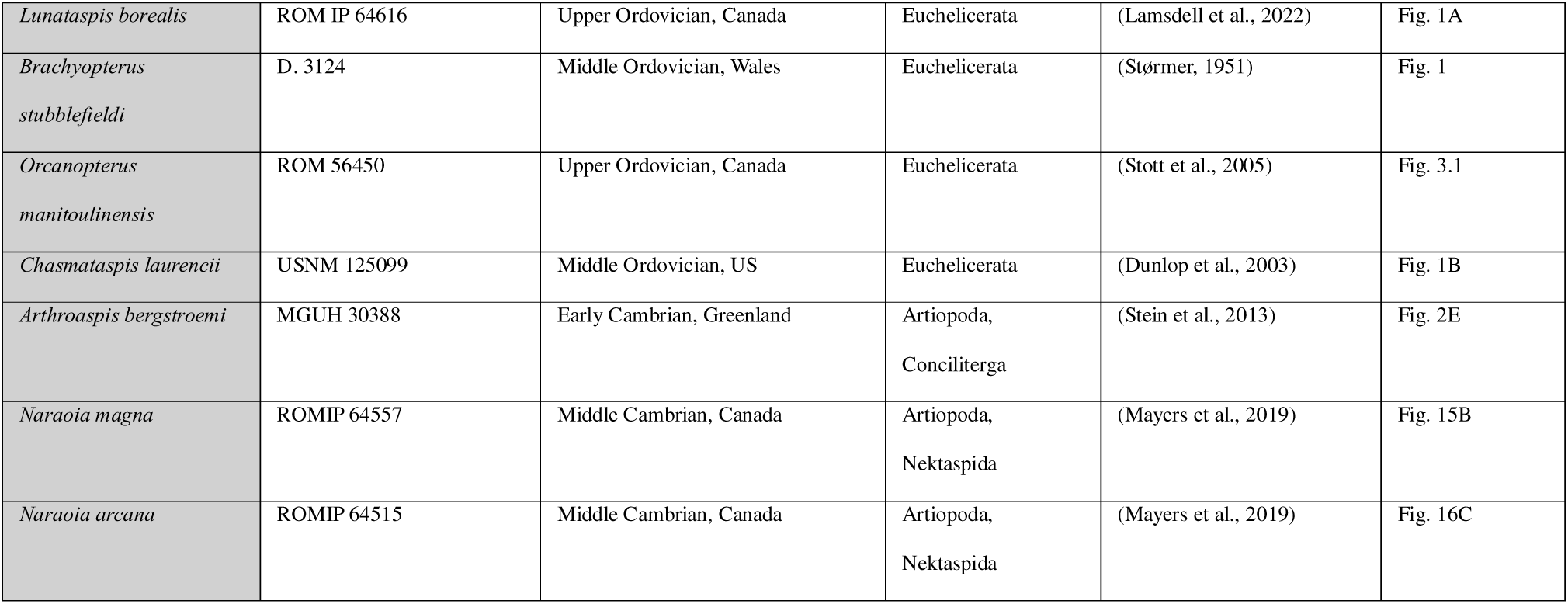
List of specimens used for the outline analyses. Figures refer to figure number in Reference column from which outline was taken.

### Material and photography

The studied material of *Zonozoe* and *Zonoscutum* is housed in the palaeontological collections of the National Museum, Prague (prefix NM L). The photographs of *Zonozoe* were taken using a Sony Alpha A7R V digital camera equipped with a Canon EF 100 mm f/2.8 Macro lens under normal lighting conditions, using a stand and a sandbox to hold the sample in place and ensure the horizontality of both the camera and the specimen. Specimens were coated with ammonium chloride prior to photography to enhance morphological details.

*Geological settings of* Zonozoe *and* Zonoscutum

The examined specimens of *Zonozoe* and *Zonoscutum* (Figure 1) come from the Ordovician deposits of the Prague Basin (Czech Republic, Barrandian area). All *Zonozoe* and *Zonoscutum* specimens represent internal moulds preserved as imprints in coarse quartzose sandstones, showing limited anatomical details. The majority of them were collected at the Děd Hill near Beroun (five specimens of *Zonozoe*; NM L 32986, NM L 23590, NM L 33029, NM L 23586, NM L 32 987) or in the fields between Trubín and Trubská villages near Králův Dvůr (one specimen of *Zonozoe* NM L 33029; see Drage et al. (2018)for details of the location), in the upper part of the Letná Formation (Chlupáč, 1965). The only known cephalon of *Zonoscutum* (NM L 33021) also originates from this latest formation and locality (Chlupáč, 1999a). Additionally, one specimen of *Zonozoe* (NM L 37385) was recovered from the Řevnice Quartzite Member within the Libeň Formation near Kařez (Rak et al., 2009), and another one has been recently reported from the Řevnice Quartzite Member near Starý Plzenec (Vokáč & Grigar, 2024).

The Libeň Formation is developed in three different members – the shallow water RCevnice Quartzite, the sequence of black shales deposited in deeper sea below the wave base in anoxic conditions, and the pyroclastic material (Havlíček, 1998). The Letná Formation is generally composed of alternating quartzose sandstones, greywackes, siltstones, and occasional shales, with sandstones representing a shallow-water environment (Chlupác & Kukal, 1988; Kraft et al., 2023).

*Zonozoe* and *Zonoscutum* have so far been discovered only in the shallow water facies of Libeň and Letná Formations. Both Letná and Libeň Formations are of Sandbian age (Upper Ordovician) (Kraft et al., 2023). From the palaeogeographical perspective, these formations within the Prague Basin were deposited in high-latitude area near the western margin of Gondwana (Kraft et al., 2023; Scotese, 2023).

### Silhouette drawing, Principal component analysis (PCA), linear discriminant analysis (LDA)

We used photographs of selected specimens to make black silhouettes of carapace outlines on a white background for selected specimens. Photographs were either taken by the authors in museum collections or sourced from the literature (see Table□1 and Figure 1). Using Adobe Illustrator 2021, we manually traced each specimen’s outline over the photograph, then exported the silhouette as a JPG file (Figure 3A). These image files served as input for the Momocs package in R (Bonhomme et al., 2014; RCoreTeam, 2024), which we used to perform the subsequent outline analyses. JPG files were imported and converted to outlines, scaled, centered, and sampled at 64-point resolution, and subjected to elliptical Fourier analysis. The function calibrate_harmonicpower_efourier was used to determine the number of harmonics required for 99.9% of the harmonic power. Elliptical Fourier coefficients were visualised using Principal Component Analysis. Linear Discriminant Analysis was used to model differences between the major groupings in the dataset (Artiopoda (non-Vicissicaudata), Euchelicerata, Vicissicaudata, *Zonozoe, Zonoscutum*). R code used for the analysis, and silhouettes in .jpg format, are supplied in Supplementary Information as Additional file 1 and 2 respectively.

**Figure 3.**
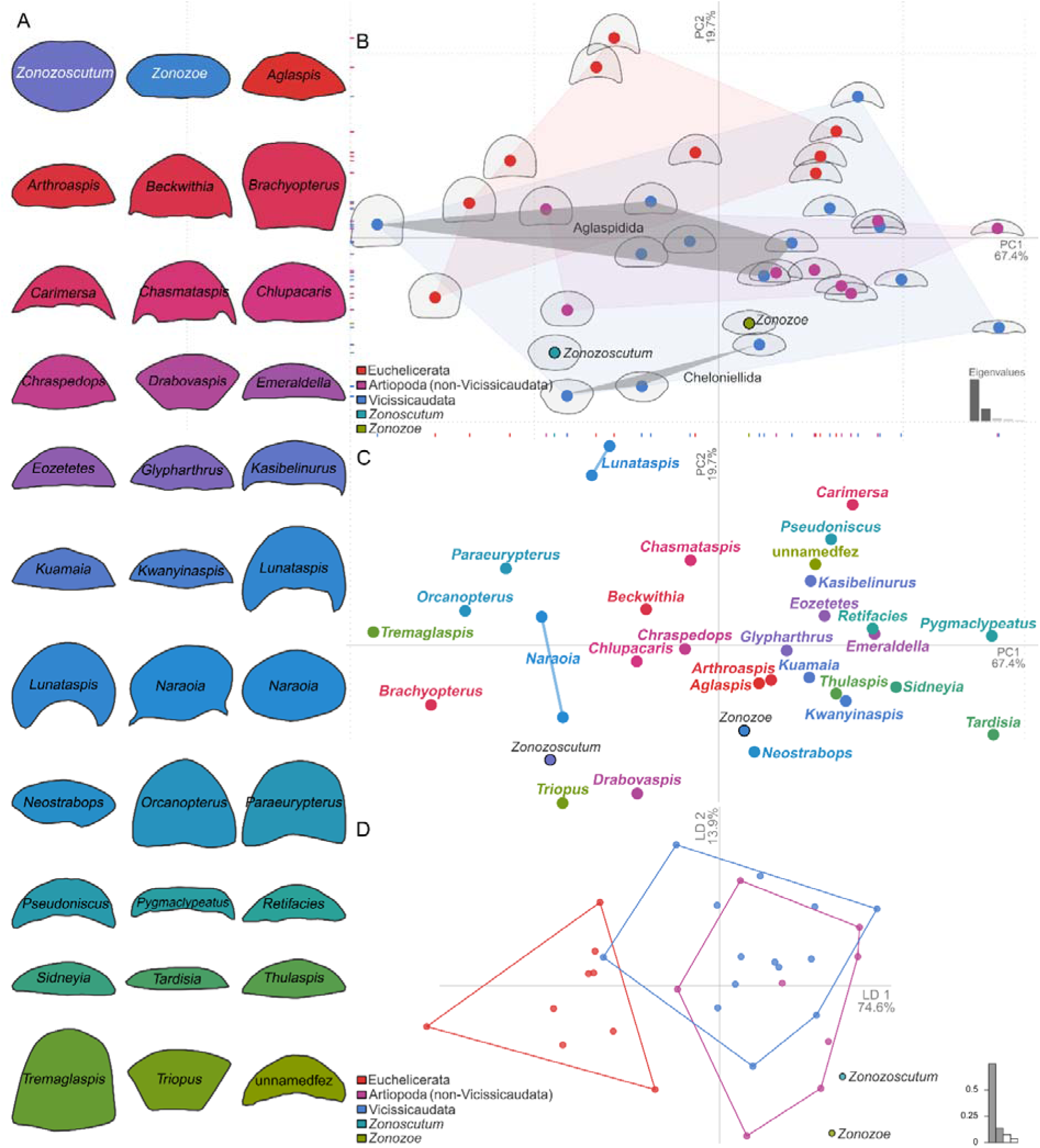
Data and Results of Principal Component Analysis and Linear Discriminant Analysis of elliptical Fourier coefficients. A, Cephalic outline of *Zonozoe drabowiensis*, *Zonoscutum solum* and the other specimens involved in the study; colors of silhouettes match colors for points in panel C. B, Position of *Zonoscutum* and *Zonozoe* in relation to major taxonomic groupings: Euchelicerata, Artiopoda (not including Vicissicaudata), and Vicissicaudata. Grey convex hulls show subgroups within Vicissicaudata, specifically Aglaspidida and Cheloniellida. Coloured marks along x and y axis indicate PC1 and PC2 values of individual specimens plotted in the space, with positions of specimens shown by filled circles. Grey outlines show variation in cephalic outline across PC1 and PC2. Histogram in bottom right shows amount of variation explained by PC1, PC2, and PC3-5; C, results by individual outline of PCA analyses. Abbreviation ‘unnamedfez’ refers to an unnamed Xiphosura from the Fezouata biota (Van Roy et al., 2010; Lamsdell & Ocon, 2025); D, Position of *Zonoscutum* and *Zonozoe* in relation to major taxonomic groupings: Euchelicerata, Artiopoda (not including Vicissicaudata), and Vicissicaudata in Linear Discriminant space. Coloured circles show positions of individual specimens. Histograms in B and D the shows the proportion of the trace for each principal component (B, C) or linear discriminant (D).

## RESULTS

### EFA, Principal component analysis (PCA), linear discriminant analysis (LDA)

Fifteen harmonics were retained, providing 99.9% of the harmonic power for the dataset. Principal component analysis reduced the dimensionality of elliptical Fourier analysis results, with PC1 and PC2 explaining 87.1% of the variation (67.4% and 19.7% respectively). Higher principal components (PC3–PC26) explain <5% of the variation each, and <15% overall. All groups overlap at least to some extent in the PCA space: Euchelicerata, Vicissicaudata, and Non□Vicissicaudata artiopodans (Figure 3B). Euchelicerates cluster toward the upper left, while non□Vicissicaudatan artiopodans occupy the area close to the origin. Vicissicaudata occupy a broader area in the morphospace, including most of the area occupied by non-Vicissicaudata, as well as areas more positive in PC1 and more negative in both PC1 and PC2. Within Vicissicaudata, aglaspidids fall near the origin of morphospace, except for *Tremaglaspis unite* (Fortey & Rushton, 2003), which plots with a negative PC1 value (Figures□3B and C). Cheloniellids form a tight cluster with negative PC2 values. Both *Zonozoe* and *Zonoscutum* lie within the lower region of the vicissicaudatan space. *Zonozoe* plots between the cheloniellid *Neostrabops martini* (Caster & Macke, 1952) and the aglaspidid *Aglaspis barrandei* (Hall, 1862)(Figures□3B and C), whereas *Zonoscutum* lies close to the cheloniellids *Triopus* and *Drabovaspis complexa* (Barrande, 1872) and the artiopodan *Naraoia* (Figures□3B and C).

Linear discriminant analysis is dominated by LD1 (74.6% of the trace), with LD2 (13.9%), with remaining LDs providing only a relatively small amount of information. Our LDA (Figure□3D) separates Euchelicerata from all other taxa but cannot separate vicissicaudates from non-vicissicaudate artiopodans (Table 2). Leave-one-out cross-validation was 48.4% (15/31) for the whole dataset, but 77.8% (7/9) for Euchelicerata. Non□Vicissicaudata artiopodans fall into the vicissicaudatans space, with leave-one-out cross-validation poor for both these groups, with most incorrect assignments placing vicissicaudatans in the non-vicissicaudatan space and vice versa (Table 2). Both *Zonozoe* and *Zonoscutum* plot at the lower right part of the LDA space, outside of all other groups but close to the Vicissicaudata/non-Vicissicaudata areas, far from the Euchelicerata area (Figure 3D).

**Table 2.**
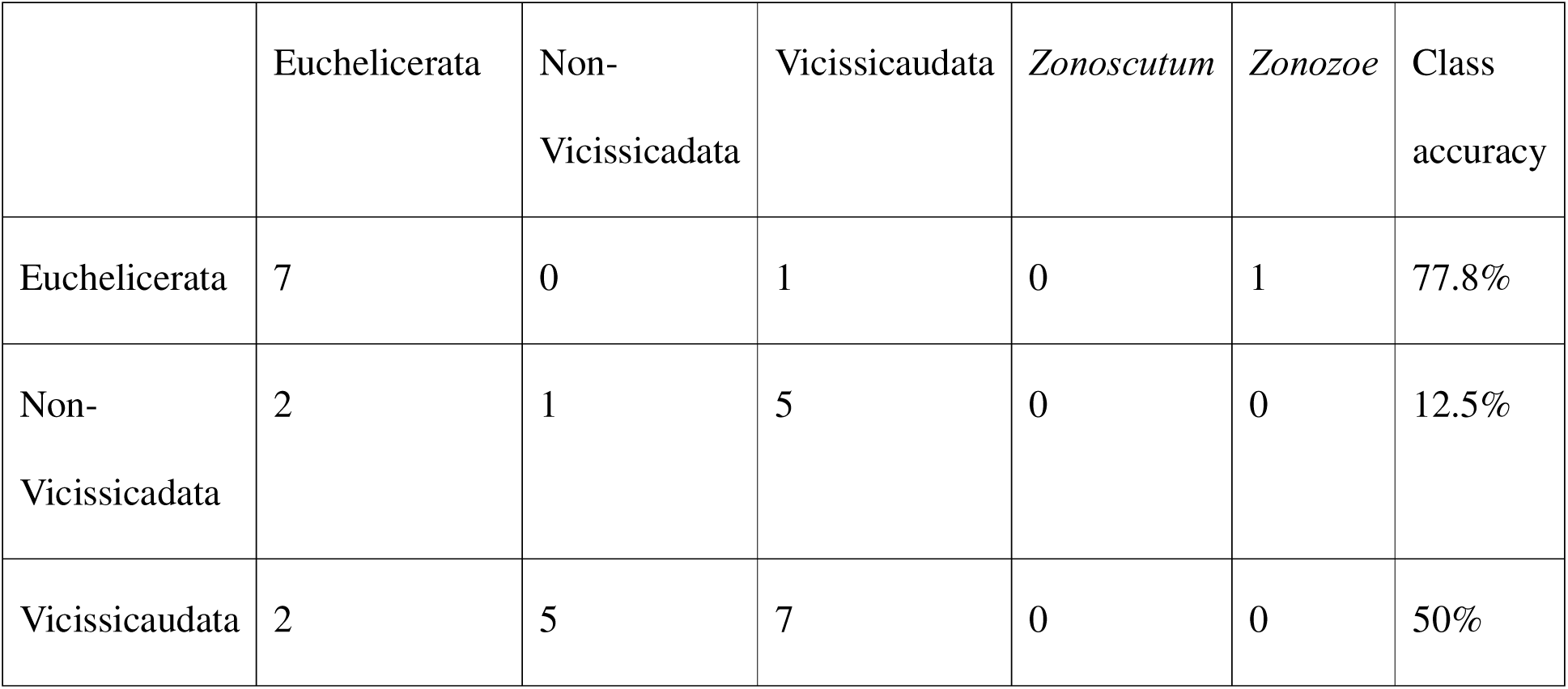
Cross validation table following Linear Discriminant Analysis, showing the classification of groups and the class accuracy. Class accuracy and classification not displayed for *Zonoscutum* and *Zonozoe* as they are represented only by a single specimen.

## DISCUSSION

Our outline analyses provide some morphological support for an artiopodan affinity for both *Zonozoe and Zonoscutum*. This was the recently lowest taxonomic level assigned to this species through the weak morphological traits with taxonomic relevance available. The PCA does not explicitly support an affinity with Aglaspidida, particularly in the case of *Zonoscutum*. Instead, both taxa are placed closer to cheloniellids than to other groups, in an intermediated position towards aglaspidids and some non-vicissicaudatan artiopodans (Figures□3B and C).

Although our analysis places *Zonozoe and Zonoscutum* near the cheloniellid cluster, we interpret these results cautiously. Key differences in cephalic morphology, which are most apparent in three-dimensional views, are not fully captured by our two-dimensional dataset, and both *Zonoscutum* and cheloniellids follow in this case. They show, respectively, a highly vaulted and extremely flattened cephalon, which may have been more affected by this methodological constraint, respectively to what was retrieved for *Zonozoe* and the other involved taxa. We believe this is the explanation behind the results of our PCA, concerning *Zonoscutum* clustering closer to the well-defined cheloniellids *Triopus* and *Drabovaspis* than to any other taxa. Based on this, we consider our method less effective for taxa with extremely vaulted or flattened cephala, such as cheloniellids and *Zonoscutum*. This limitation might have been addressed by using 3D volume data, but such data are difficult to obtain for fossils held in collections across multiple continents, and will also be impacted by flattening during or after fossilisation. Thus, we do not consider the results of our analyses strong enough to question (Van Roy et al., 2022), who rejected a close relationship between *Zonozoe*, *Zonoscutum*, and cheloniellids, especially with the coeval and sympatric *Triopus*. On the other hand, our analyses show low support for the alternative hypothesis of euchelicerate affinities, as euchelicerates are separated in the PC space from the other groups, with *Zonozoe* and *Zonoscutum* plotting far from euchelicerates and within the Vicissicaudata area.

The closest non-cheloniellid arthropod to *Zonoscutum,* is *Naroia*, followed by the euchelicerate *Brachyopterus* and the aglaspidid *A. barrandei*. Those results do not allow us to state further interpretations for *Zonoscutum*. The results for *Zonozoe,* instead, present some clearer insight. The closest taxa in our PCA to *Zonozoe* are the cheloniellid *Neostrabops* and the aglaspidid *A. barrandei*. Ortega-Hernández et al. (2013) assigned *Neostrabops* as the less derived Cheloniellida, on the basis of a shared “presence of rounded genal angles, the anterior boundaries of the trunk reflexed anterolaterally and a reduced head shield”. However, the absence of a well-developed thoracic axis, already led Van Roy (2006) to suggest it may represent the less derived form of the group. Other studies (e.g., Legg et al., 2013; Lerosey-Aubril, Zhu, et al., 2017) also placed *Neostrabops* as a basal cheloniellid, but again relied on limited synapomorphies. The lack of preservation of key traits, such as furcal rami or the morphology of terminal tergites, hinders reliable comparison with better-known forms like *Triopus* or *Cheloniellon calmani* Broili, 1932. The proximity of *Zonozoe* and *Neostrabops* in our PCA may therefore reflect more on the uncertain taxonomy of *Neostrabops* than on *Zonozoe* itself. Both fall within the vicissicaudatan morphospace, closer to aglaspidids than to other cheloniellids. Based on our results, we suggest that the taxonomical assignment of *Neostrabops* could be reconsidered towards Aglaspidida after new investigations or, in a more parsimonious way, towards Xenopoda.

On the other hand, the close PCA placement of *Zonozoe* (and *Neostrabops*) with *A. barrandei* supports a possible affinity with aglaspidids. While we do not draw a definitive conclusion, this result is consistent with previous observations, such as the position of the eyes and glabellar furrows behind them, which, while not strongly diagnostic (Van Roy et al., 2022), align with our morphometric findings and the exclusion of a possible euchelicerate affinity retrieved in our analyses (Figure 3).

Our LDA results, though more influenced by predefined categories, provide additional insight. Euchelicerates are distinguished from Artiopoda, however vicissicaudatans and non-vicissicaudatans cannot be clearly separated (Figure 3D; Table 2). This was an unexpected finding for our outline data, given the superficial similarity in dorsal morphology between some early euchelicerates and vicissicaudatans like aglaspidids (e.g. *P. anatoliensis* and *T. unite*). Previous authors have even suggested the affinities of euchelicerates with aglaspidids based on such similarities (Størmer L. 1955), but our LDA analyses were able to discriminate the two groups. Both taxa of interest, *Zonoscutum* and *Zonozoe,* are positioned closer to non-vicissicaudatan artiopods than to vicissicaudatans, in contrast with our PCA results, however, they are clearly separated from Euchelicerata. These results reinforce the more recent interpretation of aglaspidids, and by extension *Zonozoe* and *Zonoscutum*, as artiopodans, most likely vicissicaudatans, by clearly distinguishing them from euchelicerates.

## CONCLUSION

Our outline□based morphometric analyses provide quantitative support for placing *Zonozoe* and *Zonoscutum* within artiopodans. Although they plot closer to cheloniellids in the morphospace of two-dimensional carapace outlines, this could reflect methodological limits due to three□dimensional cephalic shape being compressed. Both PCA and LDA consistently separate Vicissicaudata, Aglaspidida, *Zonozoe* and *Zonoscutum* from euchelicerates. Within PCA space, they plot closest to vicissicaudata artiopods. Taken together, these results support a vicissicaudatan affinity for these taxa and reject an euchelicerates affinity. Overall, our method has proven to be a useful alternative for classifying arthropod fossils when discrete characters required for phylogenetic methods are absent. This is often the case with moulds, which rarely preserve fine anatomical details. While our approach is less precise than formal phylogenetic analysis, it allows for the inclusion of specimens that would otherwise be excluded from broader evolutionary or paleoecological studies. In our case, it provides evidence for the presence of non-cheloniellid vicissicaudatans in the Letná Formation, based on poorly preserved specimens that would likely be overlooked using only traditional qualitative methods.

## ACKNOWLEDGMENTS

We are thankful to the reviewers for their valuable comments and suggestions, which greatly improved this manuscript. This study was supported by the grant “Borse di studio SPI” from “Società Paleontologica Italiana” (SPI) awarded to L. Lu. in 2024. This study was also supported by the grants of PostDoctoral Research Fund to L.Lu. and L.C at Yunnan University. Y.L. is currently supported by the Yunnan Revitalization Talent Support Program (Grant number: 202401BC070012) from the Natural Science Foundation of Yunnan Province, China, and an Honorary visiting professor at the University of Leicester, UK funded by the Chinese Scholarship Council. SP acknowledges support from a NERC IRF NE/X017745/2. L.La is supported by the Czech Science Foundation grant GA 25-16610S and by the institutional support RVO 67985831 of the Institute of Geology of the Czech Academy of Sciences. J.B. was financially supported by Ministry of Culture of the Czech Republic through the DKRVO 2024–2028/2.III.b, National Museum, 00023272. M.N. was supported by internal grant of the Czech Geological Survey no. 311 630, which contributes to the Strategic Research Plan of the Czech Geological Survey (DKRVO/CGS 2023-2027). This paper is a contribution to IGCP project 735 “Rocks and the Rise of Ordovician Life: Filling knowledge gaps in the Early Palaeozoic Biodiversification.

## AUTHOR CONTRIBUTIONS

L.Lu. conceptualized the original idea and wrote the first draft of the manuscript. L.Lu., L.La. and L.C. collected the data. S.P. analysed the data. L.La. and L.C., and S.P. prepared the figures and wrote sections of the material and methods, results, and discussion. Y.L., J.B. and M.N. were involved in the discussion and final editing of the manuscript. All authors read and approved the final version of the manuscript.

